# Prevention of heart failure in hypertension – disentangling the role of evolving left ventricular hypertrophy and blood pressure lowering: the ALLHAT study

**DOI:** 10.1101/514323

**Authors:** Kyle Johnson, Suzanne Oparil, Barry R. Davis, Larisa G. Tereshchenko

## Abstract

**Background:** Hypertension (HTN) is a known risk factor for heart failure (HF), possibly via the mechanism of cardiac remodeling and left ventricular hypertrophy (LVH). We studied how much blood pressure (BP) change and evolving LVH contribute to the effect that lisinopril, doxazosin, amlodipine have on HF compared to chlorthalidone.

**Methods:** We conducted causal mediation analysis of Antihypertensive and Lipid-Lowering Treatment to Prevent Heart Attack Trial (ALLHAT) data. ALLHAT participants with available serial ECGs and BP measurements were included (n=29,892; mean age 67±4 y; 32% black; 56% men): 11,008 were randomized to chlorthalidone, 5,967 – to doxazosin, 6,593 – to amlodipine, and 6,324 – to lisinopril. Evolving ECG-LVH, and BP-lowering served as mediators. Incident symptomatic HF was the primary outcome. Linear regression (for mediator) and logistic regression (for outcome) models were adjusted for mediator-outcome confounders (demographic and clinical characteristics known to be associated both with both LVH/HTN and HF).

**Results:** A large majority of participants (96%) had ECG-LVH status unchanged; 4% developed evolving ECG-LVH. On average, BP decreased by 11/7 mmHg. In adjusted Cox regression analyses, progressing ECG-LVH [HR 1.78(1.43-2.22)], resolving ECG-LVH [HR 1.33(1.03-1.70)], and baseline ECG-LVH [1.17(1.04-1.31)] carried risk of incident HF. After full adjustment, evolving ECG-LVH mediated 4% of the effect of doxazosin on HF. Systolic BP-lowering mediated 12% of the effect of doxazosin, and diastolic BP-lowering mediated 10% effect of doxazosin, 7% effect of amlodipine, and borderline 9% effect of lisinopril on HF.

**Conclusions:** Evolving ECG-LVH and BP change account for 4-13% of the mechanism by which antihypertensive medications prevent HF.

## Introduction

Hypertension (HTN) is a major risk factor for heart failure (HF).^1^ HTN triggers cardiac remodeling and development of left ventricular hypertrophy (LVH), leading to subclinical organ damage, which evolves to clinically manifest HF, and ultimately, death^2^. The beneficial effect of antihypertensive treatment on HF risk is well-known,^3^ and reflected in the 2017 ACC/AHA Guideline for the Prevention, Detection, Evaluation, and Management of High Blood Pressure in Adults.^4^ HTN treatment is associated with an approximately 20-25% reduction in risk of incident HF^5^.

The Antihypertensive and Lipid-Lowering Treatment to Prevent Heart Attack Trial (ALLHAT)^6^ was a multicenter, randomized, double-blind, active-controlled trial designed to compare cardiovascular (CV) outcomes in high-risk antihypertensive patients assigned to the angiotensin-converting enzyme inhibitor (ACEi) lisinopril, the calcium channel blocker (CCB) amlodipine, and the ά-blocker doxazosin, in comparison to a thiazide-type diuretic (chlorthalidone). Incident HF was a pre-specified ALLHAT outcome. The rationale for the ALLHAT hypothesis was based on the previous demonstrations that ACEIs and CCBs are more effective than diuretics in reducing left ventricular mass index, measured by echocardiography.^7^ Contrary to expectations, the ALLHAT showed that chlorthalidone was superior to amlodipine, lisinopril, and doxazosin in preventing HF.^8, 9^ The subsequent ALLHAT HF validation study reinforced original ALLHAT results.^10–12^

While ALLHAT answered question about the comparative effectiveness of antihypertensive treatments for HF prevention, mechanisms behind this HF prevention remain incompletely understood. The extent to which the effect of a CCB, ACEi, and an ά-blocker (as compared to a diuretic) on incident HF is mediated by evolving LVH and blood pressure (BP) lowering *per se* remains unknown. This study aimed to quantify the extent to which the effect of lisinopril, amlodipine, and doxazosin (as compared to chlorthalidone) on incident HF is mediated by evolving LVH and BP lowering. We hypothesized that evolving ECG LVH and BP lowering are mechanisms behind previously observed differences in the rate of incident HF in hypertensive ALLHAT participants randomized to lisinopril, amlodipine, and doxazosin, in comparison to those randomized to chlorthalidone.

## Methods

For this study, we used the ALLHAT dataset, publicly available from the National Heart, Lung, and Blood Institute, via BioLINCC. The study was reviewed by an Oregon Health and Science University Institutional Review Board and determined that it did not require further review due to the de-identified nature of publicly available dataset.

### Study population

The ALLHAT design and rationale have been described previously.^6^ Briefly, ALLHAT enrolled adults age 55 and above, with HTN and at least one risk factor [documented coronary heart disease (CHD), type II diabetes mellitus, LVH on ECG or echocardiogram, smoking, high-density lipoprotein (HDL) < 35mg/dL, or ST-T ECG changes indicative of ischemia]. Symptomatic HF patients or those with LVEF <35%, patients with recent myocardial infarction (MI), stroke, or poorly controlled HTN were excluded.

In this study, we included ALLHAT participants with available assessment of evolving LVH status, and dynamic BP changes. We excluded participants with missing covariates. Final study population included 29,892 participants: 11,008 were randomized to chlorthalidone, 5,967 – to doxazosin, 6,593 – to amlodipine, and 6,324 – to lisinopril (Figure 1).

**Figure 1.**
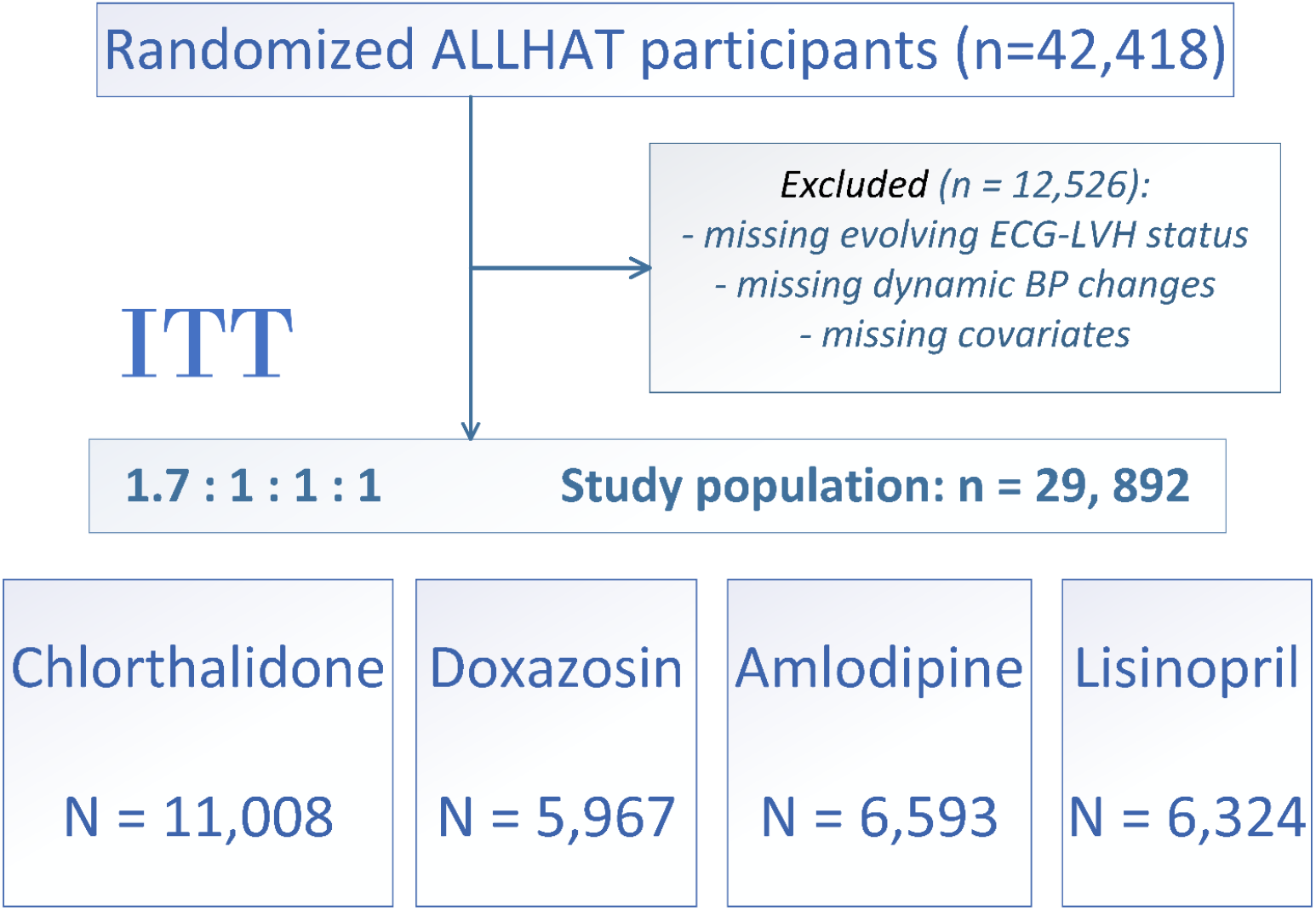
Flow diagram of exclusion criteria applied to achieve the final study population for this secondary analysis of Antihypertensive and Lipid-Lowering Treatment to Prevent Heart Attack Trial (ALLHAT) data.

### ECG analysis: Evolving LVH during follow-up

ECGs were recorded at the study sites at baseline and biannually during follow-up. Minnesota coding^13^ of serial ECG changes (Table 1) was performed in the ECG core center at the University of Minnesota in Minneapolis by reviewers who were blinded to treatment assignments. Minnesota codes *3-1* and *3-3* are high left R amplitude patterns (relevant to LVH) as measured on the next to last complete normal beat. Code *3-1* was coded if any of the following 3 criteria are present: (1) R amplitude >26 mm in either lead V5 or V6; (2) R amplitude >20 mm in any of leads I, II, III, or aVF; (3) R amplitude >12 mm in lead aVL. Code *3-3* was coded if one or both of the following two criteria is present: (1) R-wave amplitude >15 mm but ≤20 mm in lead I; R-wave amplitude in V5 or V6 plus S or QS amplitude in V1 >35 mm.

**Table 1.**
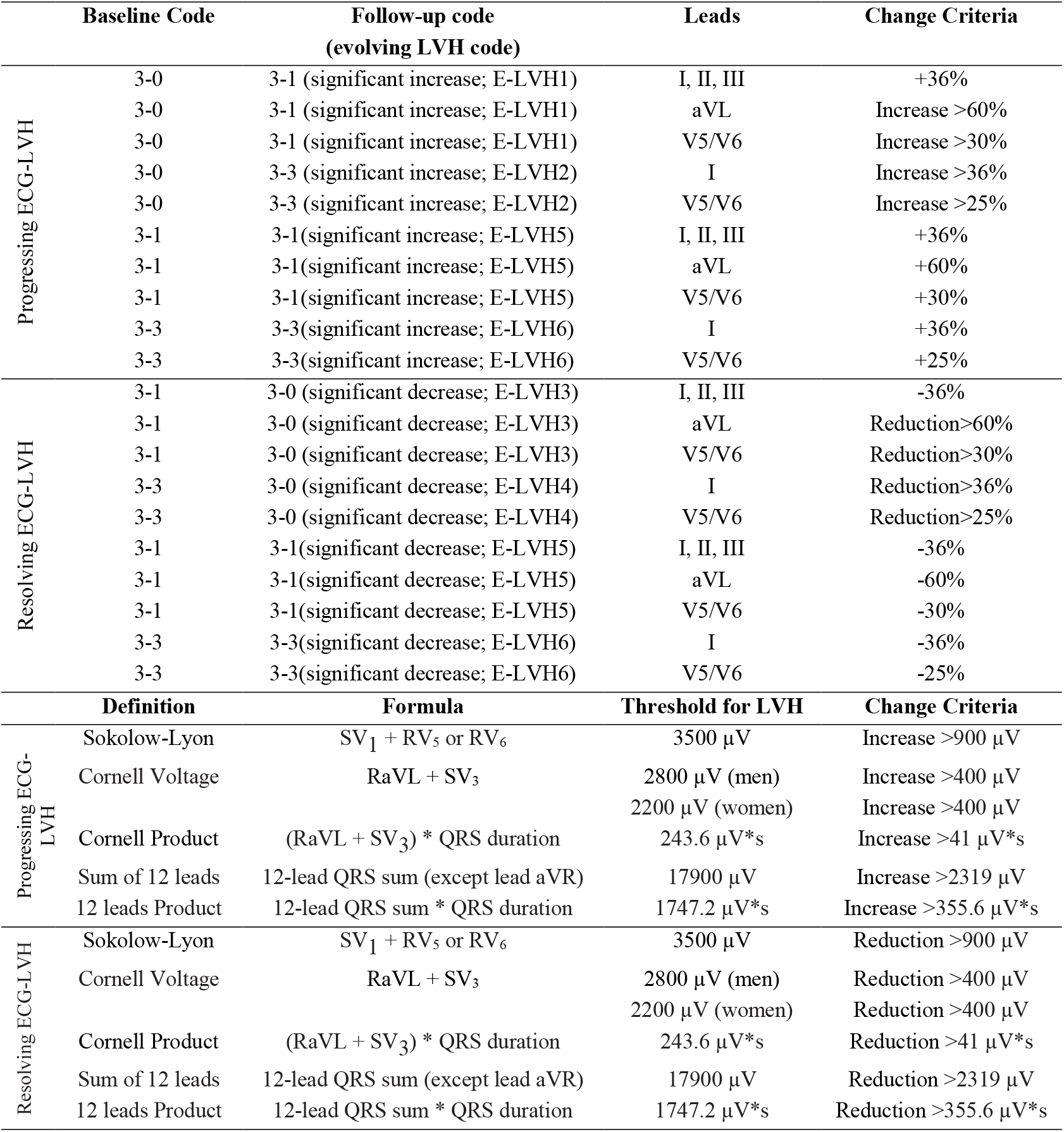
Minnesota Code definitions of evolving ECG-LVH^13^

Serial ECG changes were assessed during follow-up, which required at least two ECGs. LVH was examined in clinic (non-hospital) ECG recordings. The Minnesota Code allowed for objective classification of evolving LVH over time by setting limits to the percentage of change in voltage that occurs in serial ECGs (Table 1). At the first step, it was determined in which lead the most severe 3-code occurred. Code *3-1* was considered more severe than Code *3-3*. If both ECGs had the same 3-code, the follow-up record determined which lead to use to compare with the reference ECG. If the 3-code occurred in different leads, the following hierarchy was used to determine which lead to compare: V5 /V6 (whichever R-amplitude is higher)>I>II>III> aVL.

Evolving LVH (Table 1) was coded as either significant progression (including newly diagnosed ECG-LVH), or significant resolution (including complete resolution of ECG-LVH). In addition, several ECG-LVH definitions were included (Sokolow-Lyon, Cornell Voltage, Cornell Product, Sum of 12 leads, 12 leads Product). Table 1 reports thresholds that were used to define evolving ECG-LVH.^13^

### Blood pressure changes during the course of the trial

BP was measured at every follow-up visit (every 3 months for the 1^st^ year and every 4 months thereafter. At each visit, BP was recorded as an average of two measurements. To calculate achieved BP lowering during the trial, we subtracted baseline BP from the BP obtained at the latest in-trial study visit available at year 1, 2, 3, 4, 5, or 6 from baseline, thus obtaining estimates of the *‘greatest’* BP control. In addition, we conducted sensitivity analyses with three other definitions of BP lowering. By subtracting baseline BP from the BP obtained at the next in-trial study visit available, we obtained estimates of the ‘*fastest’* BP control. We also divided the greatest and fastest BP control estimates by the baseline BP, obtaining relative greatest and fastest BP lowering.

### Primary outcome: Incident heart failure

Incident symptomatic congestive HF as defined by the ALLHAT investigators was a primary outcome in this study. Diagnosis of symptomatic congestive HF required the presence of both:

(1) Paroxysmal nocturnal dyspnea, or dyspnea at rest, or New York Heart Association class III symptomps, or orthopnea, and (2) rales, or ankle edema (2+ or greater), or sinus tachycardia of 120 beats/minute or more after 5 minutes at rest, or cardiomegaly by chest X-ray, or chest X-ray characteristic of congestive HF, or S3 gallop, or jugular venous distention. The incident HF outcome was validated by the ALLHAT HF validation study.^10^ In the current study, hospitalized / fatal HF was included as a secondary outcome.

### Covariates

Baseline BP was calculated as an average of two BP determinations taken at least one day apart, with each determination being an average of 2 measurements.

Baseline ECG-LVH was based on any ECG within the past 2 years. Baseline ECG-LVH definition included any one of the following: (1) R amplitude in V5 or V6 > 26 mm, (2) R amplitude in V5 or V6 plus S amplitude in V1 > 35 mm, (3) R amplitude in aVL > 12 mm, (4) R amplitude in Lead I > 15 mm, (5) R amplitude in Leads II or III, or aVF > 20 mm, (6) R amplitude in Lead I plus S amplitude in Lead III > 25 mm, (7) R amplitude in aVL plus S amplitude in V3 > 28 mm for men or > 22 mm for women, or (8) computerized ECG machine documented LVH.

Echocardiographic LVH (Echo-LVH) was defined as combined wall (posterior wall plus interventricular septum) thickness ≥ 25 mm on any echocardiogram in the past 2 years.

Baseline medical history was determined by the study investigators by a combination of chart review and questioning during a routine office visit. HTN history determined whether participants were treated for at least 2 months, were treated for less than 2 months, or were untreated. History of MI or stroke was at least 6 months old. History of revascularization included history of angioplasty, stenting, atherectomy, bypass surgery [coronary; peripheral vascular; carotid; vertebrobasilar], or aortic aneurysm repair. Presence of major ST segment depression or T wave elevation on any ECG in the past two years was identified. History of other atherosclerotic cardiovascular disease (CVD) included documented peripheral artery disease or cerebrovascular disease. Baseline CHD history included known prior MI (including silent MI), angina, cardiac arrest, angiographically defined coronary stenosis more than 50%, reversible perfusion defects on cardiac scintigraphy, or prior coronary revascularization procedures. Type II diabetes was defined as fasting plasma glucose > 140 mg/dl [7.77 mmol/L] or non-fasting plasma glucose > 200 mg/dl [11.1 mmol/L] in the past 2 years and/or current treatment with insulin or oral hypoglycemic agents. History of HDL cholesterol < 35 mg/dl (0.91 mmol/l) on any 2 or more determinations within past 5 years was included. History of smoking was also obtained.

### Statistical analysis

All continuous variables are presented as means±standard deviation (SD). ANOVA and χ^2^ test was used for unadjusted comparison of clinical characteristics in participants with evolving ECG-LVH. To determine association of clinical characteristics with achieved in-trial BP changes, we used multivariable linear regression models, minimally adjusted for age, sex, and race/ethnicity. Intention-to-treat (ITT) randomization assignment was used for definition of antihypertensive treatment groups.

Minimally adjusted (by age, sex, and race/ethnicity) Cox regression models were used to describe associations of baseline clinical characteristics, evolving ECG-LVH, and BP-lowering with two different definitions of incident HF, for comparison. Associations between BP-lowering (continuous variable) and HF risk were also evaluated using adjusted (as above) Cox regression models incorporating cubic splines with 4 knots.

We conducted causal mediation analysis^14^, allowing for treatment-mediator interaction in the logistic regression, using counterfactual definitions of direct and indirect effects, as implemented by VanderWeele and colleagues.^15^ Two models were estimated: a linear model for the mediator conditional on treatment and covariates, and a logistic model for the outcome conditional on treatment, the mediator, and covariates. Our study design is well-suited for mediation analysis, as randomization eliminated exposure-outcome and exposure-mediator confounding. Two mediators were studied (Figure 2): (1) evolving ECG-LVH, and (2) BP lowering over the course of the trial. We adjusted for mediator-outcome confounders^11, 16^, which were measured at baseline: demographic (age, sex, race and ethnicity) and clinical characteristics known to be associated both with LVH/HTN and HF: common risk factors (body mass index [BMI], smoking, diabetes), HTN history (levels of baseline systolic BP (SBP) and diastolic BP (DBP), baseline use of antihypertensive medications, ECG- or echo-LVH), CHD or CVD history, coronary revascularization, major ST depression or T-wave inversion, HDL<35 mg/dL twice in the past 5 years, and participation in the lipid-lowering ALLHAT trial. A natural direct effect represents the influence of antihypertensive treatment that is independent of evolving ECG-LVH or BP-lowering, in the absence of evolving ECG-LVH or BP changes (e.g. via pleiotropic effects or drug-specific pharmacodynamics). A controlled direct effect represents the effect of antihypertensive drug at certain level of mediator (at progressing/resolving ECG-LVH with a reference at absent evolving ECG-LVH, and at tertiles of BP changes), allowing measurement of interaction between treatment and a mediator. A mediated effect represents the influence of antihypertensive drug that can be explained by its influence on evolving ECG-LVH or dynamic BP changes achieved over the course of the trial. To assess the extent of mediation, we estimated the proportion mediated as a ratio of DE*(ME-1)/(DE*ME-1), where DE is direct effect and ME is mediated effect.

**Figure 2.**
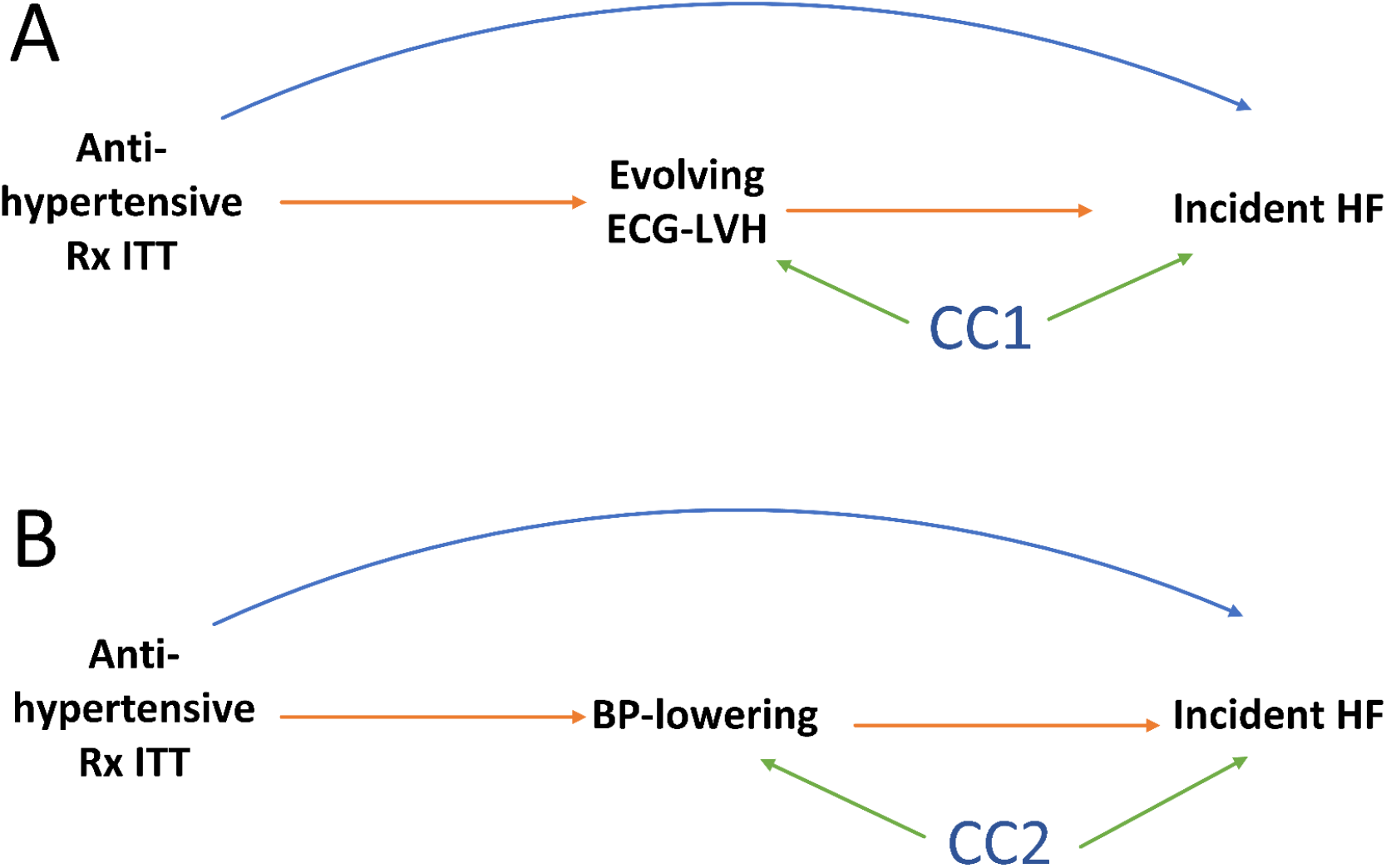
Directed acyclic graph to illustrate possible structural relationships between randomized antihypertensive treatment (Rx) in intention-to-treat (ITT) analysis, evolving ECG-LVH (**A**) or BP-lowering (**B**), and incident HF. CC denotes common causes (confounding factors), measured and unmeasured. The mediated effect is represented by the pathway from antihypertensive Rx to incident HF that goes through (**A**) evolving ECG-LVH or (**B**) BP-lowering. The direct effect is the pathway from antihypertensive Rx straight to incident HF.

#### Sensitivity analyses

To test robustness of our findings, we repeated analyses with different definitions of BP lowering, expressed as: (1) fastest BP control; (2) relative greatest BP control; (3) relative fastest BP control.

Statistical analyses were performed using STATA MP 15.1 (StataCorp LLC, College Station, TX). Given the many multivariate and interaction analyses performed, statistical significance at the 0.05 level should be interpreted cautiously.

## Results

### Study population

Study population (Table 2) was identical to previously reported ALLHAT population,^8, 9^ maintaining treatment groups randomization ratio 1.7:1:1:1. After median 3.1 years follow-up in doxazosin group, and 5.0 years in other 3 groups, there were 2,049 incident HF outcomes, including 1,598 hospitalized/fatal HF outcomes.

**Table 2.**
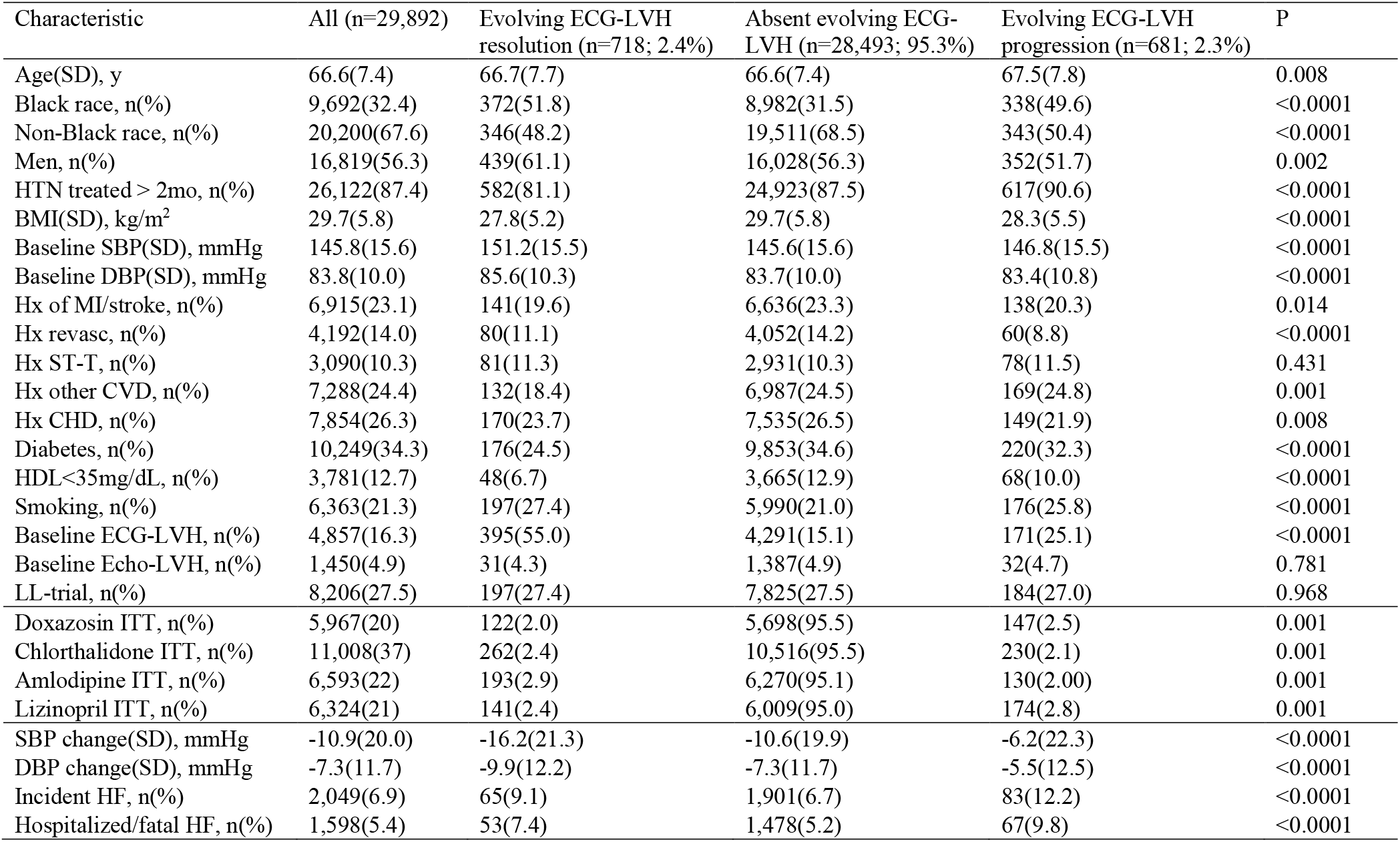
Clinical characteristics of study participants with evolving ECG-LVH increase or decrease

| Characteristic | All (n=29,892) | Evolving ECG-LVH resolution (n=718; 2.4%) | Absent evolving ECG-LVH (n=28,493; 95.3%) | Evolving ECG-LVH progression (n=681; 2.3%) | P |
| --- | --- | --- | --- | --- | --- |
| Age(SD), y | 66.6(7.4) | 66.7(7.7) | 66.6(7.4) | 67.5(7.8) | 0.008 |
| Black race, n(%) | 9,692(32.4) | 372(51.8) | 8,982(31.5) | 338(49.6) | <0.0001 |
| Non-Black race, n(%) | 20,200(67.6) | 346(48.2) | 19,511(68.5) | 343(50.4) | <0.0001 |
| Men, n(%) | 16,819(56.3) | 439(61.1) | 16,028(56.3) | 352(51.7) | 0.002 |
| HTN treated > 2mo, n(%) | 26,122(87.4) | 582(81.1) | 24,923(87.5) | 617(90.6) | <0.0001 |
| BMI(SD), kg/m<sup>2</sup> | 29.7(5.8) | 27.8(5.2) | 29.7(5.8) | 28.3(5.5) | <0.0001 |
| Baseline SBP(SD), mmHg | 145.8(15.6) | 151.2(15.5) | 145.6(15.6) | 146.8(15.5) | <0.0001 |
| Baseline DBP(SD), mmHg | 83.8(10.0) | 85.6(10.3) | 83.7(10.0) | 83.4(10.8) | <0.0001 |
| Hx of MI/stroke, n(%) | 6,915(23.1) | 141(19.6) | 6,636(23.3) | 138(20.3) | 0.014 |
| Hx revasc, n(%) | 4,192(14.0) | 80(11.1) | 4,052(14.2) | 60(8.8) | <0.0001 |
| Hx ST-T, n(%) | 3,090(10.3) | 81(11.3) | 2,931(10.3) | 78(11.5) | 0.431 |
| Hx other CVD, n(%) | 7,288(24.4) | 132(18.4) | 6,987(24.5) | 169(24.8) | 0.001 |
| Hx CHD, n(%) | 7,854(26.3) | 170(23.7) | 7,535(26.5) | 149(21.9) | 0.008 |
| Diabetes, n(%) | 10,249(34.3) | 176(24.5) | 9,853(34.6) | 220(32.3) | <0.0001 |
| HDL<35mg/dL, n(%) | 3,781(12.7) | 48(6.7) | 3,665(12.9) | 68(10.0) | <0.0001 |
| Smoking, n(%) | 6,363(21.3) | 197(27.4) | 5,990(21.0) | 176(25.8) | <0.0001 |
| Baseline ECG-LVH, n(%) | 4,857(16.3) | 395(55.0) | 4,291(15.1) | 171(25.1) | <0.0001 |
| Baseline Echo-LVH, n(%) | 1,450(4.9) | 31(4.3) | 1,387(4.9) | 32(4.7) | 0.781 |
| LL-trial, n(%) | 8,206(27.5) | 197(27.4) | 7,825(27.5) | 184(27.0) | 0.968 |
| Doxazosin ITT, n(%) | 5,967(20) | 122(2.0) | 5,698(95.5) | 147(2.5) | 0.001 |
| Chlorthalidone ITT, n(%) | 11,008(37) | 262(2.4) | 10,516(95.5) | 230(2.1) | 0.001 |
| Amlodipine ITT, n(%) | 6,593(22) | 193(2.9) | 6,270(95.1) | 130(2.00) | 0.001 |
| Lizinopril ITT, n(%) | 6,324(21) | 141(2.4) | 6,009(95.0) | 174(2.8) | 0.001 |
| SBP change(SD), mmHg | -10.9(20.0) | -16.2(21.3) | -10.6(19.9) | -6.2(22.3) | <0.0001 |
| DBP change(SD), mmHg | -7.3(11.7) | -9.9(12.2) | -7.3(11.7) | -5.5(12.5) | <0.0001 |
| Incident HF, n(%) | 2,049(6.9) | 65(9.1) | 1,901(6.7) | 83(12.2) | <0.0001 |
| Hospitalized/fatal HF, n(%) | 1,598(5.4) | 53(7.4) | 1,478(5.2) | 67(9.8) | <0.0001 |

### Serial ECG changes: evolving ECG-LVH

Overall, 58,366 serial ECG changes were evaluated. ECG-LVH resolution was observed in about 2% of participants, and in another 2% ECG-LVH progressed (Table 2). The majority of participants had no evolving ECG-LVH changes. ALLHAT participants with evolving ECG-LVH were more likely black males, current smokers with lower BMI, but less likely having CHD/MI history. As expected, baseline ECG-LVH was more frequent in participants with resolving ECG-LVH. Baseline LVH by echocardiogram was similar in all 3 groups, and was very infrequent (4-5%). Participants with resolving LVH by ECG were more likely diabetic, less likely to have been treated before the onset of the trial, and achieved the greatest degree of BP-lowering in-trial. Incident HF was significantly more frequent in participants with evolving ECG-LVH (Table 2). Doxazosin and lisinopril ITT were more likely to be associated with progressing ECG-LVH, and less likely associated with ECG-LVH reduction. In contrast, chlorthalidone and amlodipine ITT were more likely to be associated with ECG-LVH reduction, and less likely associated with ECG-LVH progression (Table 2).

### Dynamic changes in Blood Pressure in-trial

The first (Q1), second (Q2), and third (Q3) tertiles of the greatest BP-lowering were −32/−19±10/6 mmHg, −11/−7±5/3mmHg, and +11/6±12/7 mmHg, respectively. Q1, Q2, and Q3 of the fastest BP-lowering were −28/−16±10/6 mmHg, −7/−4±5/3 mmHg, and +14/8±12/6 mmHg, accordingly. Hispanic ethnicity, previously untreated HTN, higher baseline levels of SBP/DBP (Figure 3) and baseline ECG-LVH were associated with greater SBP and DBP lowering in-trial (Table 3). In contrast, presence of diabetes was associated with a SBP increase of nearly 2 mmHg. Older age was associated with greater SBP-lowering but slight DBP-increase. History of CHD/CVD did not affect the degree of BP-lowering in-trial. Compared to chlorthalidone, doxazosin was associated with significant SBP increase (by nearly 2 mmHg), whereas amlodipine was associated with significant SBP and DBP decrease. Lisinopril was associated with greater DBP (but not SBP) lowering than chlorthalidone (Table 3). Participants in the doxazosin arm who developed HF had the greatest degree of BP-lowering (both SBP/DBP) in-trial (~6/2 mmHg lower than by diuretic), which contrasted with overall weak BP-lowering effect of doxazosin in the trial (Table 3).

**Figure 3:**
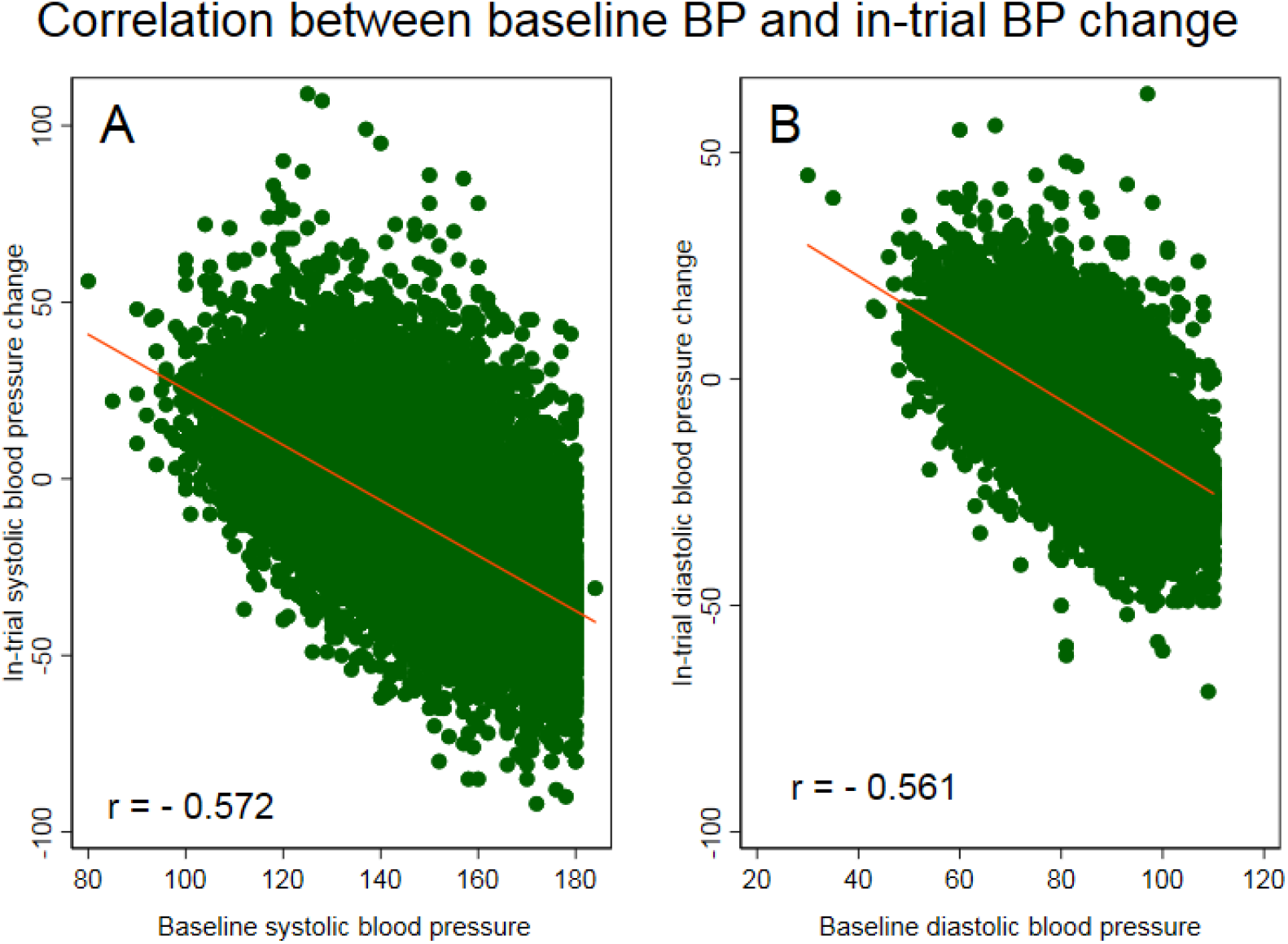
Scatterplots of (A) in-trial SBP change (Y-axis) against baseline SBP (X-axis), and (B) DBP change (Y-axis) against baseline DBP (X-axis). A line of the best linear fit is shown.

**Table 3.** Associations of clinical characteristics with BP change in-trial, in linear regression models

| Characteristic | Systolic BP change(95% CI), mmHg | P | Diastolic BP change(95% CI), mmHg | P |
| --- | --- | --- | --- | --- |
| Age, per 1 y increase | -0.13(-0.16 to -0.10) | **<0.0001** | +0.02(0.006-0.04) | **0.009** |
| Race/ethnicity: White non-hispanic | Reference |  | Reference |  |
| Black non-hispanic | +3.09(2.57-3.61) | **<0.0001** | +1.59(1.29-1.90) | **<0.0001** |
| White Hispanic | -3.64(-4.42 to -2.86) | **<0.0001** | -1.32(-1.78 to -0.87) | **<0.0001** |
| Black Hispanic | -3.79(-5.37 to -1.19) | **<0.0001** | -0.89(-1.82 to 0.03) | 0.076 |
| Women | +0.44(-0.02 to 0.90) | 0.063 | +0.67(0.40-0.94) | **<0.0001** |
| HTN treated: ≥ 2months | Reference |  | Reference |  |
| < 2 months | -6.89(-8.16 to -5.61) | **<0.0001** | -3.70(-4.45 to -2.94) | **<0.0001** |
| Not treated | -11.85(-12.61 to -11.08) | **<0.0001** | -5.74(-6.19 to -5.29) | **<0.0001** |
| BMI, per 1 kg/m<sup>2</sup> increase | +0.04(-0.0008 to 0.08) | 0.055 | -0.004(-0.03 to 0.02) | 0.701 |
| Baseline SBP, per 1 mmHg increase | -0.78(-0.80 to -0.77) | **<0.0001** | -0.29(-0.29 to -0.28) | **<0.0001** |
| Baseline DBP, per 1 mmHg increase | -0.71(-0.73 to -0.69) | **<0.0001** | -0.72(-0.73 to -0.71) | **<0.0001** |
| Hx of MI/stroke | -0.22(-0.77 to 0.32) | 0.423 | +0.22(-0.10 to 0.54) | 0.178 |
| Hx revascularization | +0.07(-0.60 to 0.74) | 0.843 | +0.08(-0.31 to 0.47) | 0.692 |
| Hx ST-T changes | -0.49(-1.24 to 0.25) | 0.194 | -0.12(-0.56 to 0.32) | 0.592 |
| Hx other CVD | -0.32(-0.85 to 0.22) | 0.245 | +0.30(-0.01 to 0.61) | 0.061 |
| Hx CHD | -0.62(-1.15 to -0.09) | **0.022** | +0.03(-0.28 to 0.34) | 0.827 |
| Diabetes | +1.64(1.17-2.11) | **<0.0001** | +0.20(-0.08 to 0.48) | 0.154 |
| HDL<35mg/dL | +0.86(0.17-1.55) | **0.014** | +0.30(-0.11 to 0.70) | 0.152 |
| Smoking: never | Reference |  | Reference |  |
| Past | -0.12(-0.66 to 0.43) | 0.675 | -0.15(-0.84 to 0.16) | 0.334 |
| Current | -1.13(-1.77 to -0.49) | **0.001** | -0.46(-0.84 to -0.09) | **0.016** |
| Baseline ECG-LVH | -1.58(-2.20 to -0.95) | **<0.0001** | -0.95(-1.31 to -0.58) | **<0.0001** |
| Baseline Echo-LVH | -0.97(-2.02 to 0.08) | 0.071 | +0.53(-0.08 to 1.15) | 0.091 |
| Treatment arm: Chlorthalidone ITT | Reference |  | Reference |  |
| Doxazosin ITT | +1.68(1.05-2.30) | **<0.0001** | -0.29(-0.66 to 0.08) | 0.124 |
| Amlodipine ITT | -1.25(-1.86 to -0.65) | **<0.0001** | -2.09(-2.44 to -1.73) | **<0.0001** |
| Lizinopril ITT | -0.16(-0.77 to 0.46) | 0.616 | -1.37(-1.73 to -1.01) | **<0.0001** |
| Incident HF | -2.66(-3.56 to -1.76) | **<0.0001** | -1.23(-1.76 to -0.70) | **<0.0001** |
| Incident HF ## Doxazosin | -5.33(-7.86 to -2.80) | **<0.0001** | -1.88(-3.36 to -0.40) | **0.013** |
| Hospitalized/Fatal HF | -2.56(-3.57 to -1.55) | **<0.0001** | -1.22(-1.81 to -0.62) | **<0.0001** |
| Hospitalized/fatal HF ## Doxazosin | -5.97(-8.85 to -3.09) | **<0.0001** | -2.06(-3.74 to 0.37) | **0.017** |

### Risk factors for Heart Failure

As expected, age, ethnicity, history of HTN, CHD, and CVD, as well as ECG-LVH were associated with increased risk of HF (Table 4). There were very little differences between risk factors of two incident HF outcomes: incident symptomatic HF and hospitalized/fatal HF.

**Table 4.** Associations of clinical characteristics with incident heart failure in Cox regression models

| Characteristic | Incident symptomatic HF HR(95%CI) | P | Hospitalized/Fatal HF HR(95%CI) | P |
| --- | --- | --- | --- | --- |
| Age, per 1 y increase | 1.06(1.05-1.06) | **<0.0001** | 1.06(1.05-1.07) | **<0.0001** |
| Race/ethnicity: White non-hispanic | Reference |  | Reference |  |
| Black non-hispanic | 0.94(0.86-1.04) | 0.234 | 0.97(0.87-1.09) | 0.634 |
| White Hispanic | 0.41(0.32-0.51) | **<0.0001** | 0.47(0.37-0.60) | **<0.0001** |
| Black Hispanic | 0.50(0.33-0.77) | **0.002** | 0.54(0.34-0.88) | **0.013** |
| Women | 0.91(0.84-0.999) | **0.048** | 0.93(0.84-1.03) | 0.177 |
| HTN treated: ≥ 2months | Reference | | Reference | |
| < 2 months | 1.04(0.81-1.34) | 0.732 | 1.21(0.93-1.58) | 0.148 |
| Not treated | 0.68(0.57-0.81) | **<0.0001** | 0.71(0.58-0.87) | **0.001** |
| BMI, per 1 kg/m<sup>2</sup> increase | 1.05(1.04-1.05) | **<0.0001** | 1.04(1.03-1.05) | **<0.0001** |
| Baseline SBP, per 1 mmHg increase | 1.006(1.004-1.009) | **<0.0001** | 1.009(1.006-1.01) | **<0.0001** |
| Baseline DBP, per 1 mmHg increase | 0.990(0.986-0.995) | **<0.0001** | 0.990(0.985-0.995) | **<0.0001** |
| Hx of MI/stroke | 1.75(1.59-1.91) | **<0.0001** | 1.78(1.61-1.98) | **<0.0001** |
| Hx revascularization | 1.73(1.55-1.92) | **<0.0001** | 1.65(1.46-1.87) | **<0.0001** |
| Hx ST-T changes | 1.10(0.96-1.26) | 0.159 | 1.14(0.98-1.33) | 0.080 |
| Hx other CVD | 1.26(1.15-1.39) | **<0.0001** | 1.27(1.14-1.41) | **<0.0001** |
| Hx CHD | 1.66(1.52-1.82) | **<0.0001** | 1.62(1.46-1.80) | **<0.0001** |
| Diabetes | 1.71(1.57-1.87) | **<0.0001** | 1.85(1.67-2.04) | **<0.0001** |
| HDL<35mg/dL | 0.98(0.86-1.11) | 0.731 | 0.97(0.84-1.13) | 0.718 |
| Smoking: never | Reference |  | Reference |  |
| Past | 1.19(1.08-1.32) | **0.001** | 1.21(1.08-1.36) | **0.001** |
| Current | 1.07(0.94-1.23) | 0.312 | 1.17(1.01-1.36) | **0.036** |
| Baseline ECG-LVH | 1.17(1.04-1.31) | **0.008** | 1.16(1.02-1.32) | **0.023** |
| Baseline Echo-LVH | 1.00(0.92-1.21) | 0.972 | 1.06(0.86-1.32) | 0.576 |
| Evolving ECG-LVH: absent | Reference |  | Reference |  |
| Resolving | 1.33(1.03-1.70) | **0.026** | 1.39(1.05-1.83) | **0.020** |
| Progressing | 1.78(1.43-2.22) | **<0.0001** | 1.84(1.44-2.35) | **<0.0001** |
| SBP lowering by 3-19 mmHg (Q2): Reference |  |  |  | **<0.0001** |
| by 20 mmHg or more (Q1) | 1.34(1.20-1.49) | **<0.0001** | 1.41(1.25-1.60) | **<0.0001** |
| By 2 mmHg or less (Q3) | 1.08(0.97-1.21) | 0.154 | 1.14(1.01-1.30) | **0.039** |
| DBP lowering by 2-11 mmHg (Q2): Reference |  |  |  |  |
| by 12 mmHg or more (Q1) | 1.31(1.18-1.45) | **<0.0001** | 1.32(1.17-1.49) | **<0.0001** |
| By 1 mmHg or less (Q3) | 1.09(0.97-1.21) | 0.155 | 1.12(0.98-1.27) | 0.088 |

Evolving ECG-LVH was associated with incident HF (Figure 4), although progressing ECG-LVH carried larger risk, as compared to resolving ECG-LVH. Evolving LVH was associated with incident HF in three out of four treatment groups (P_interaction_=0.056; Figure 5).

**Figure 4.**
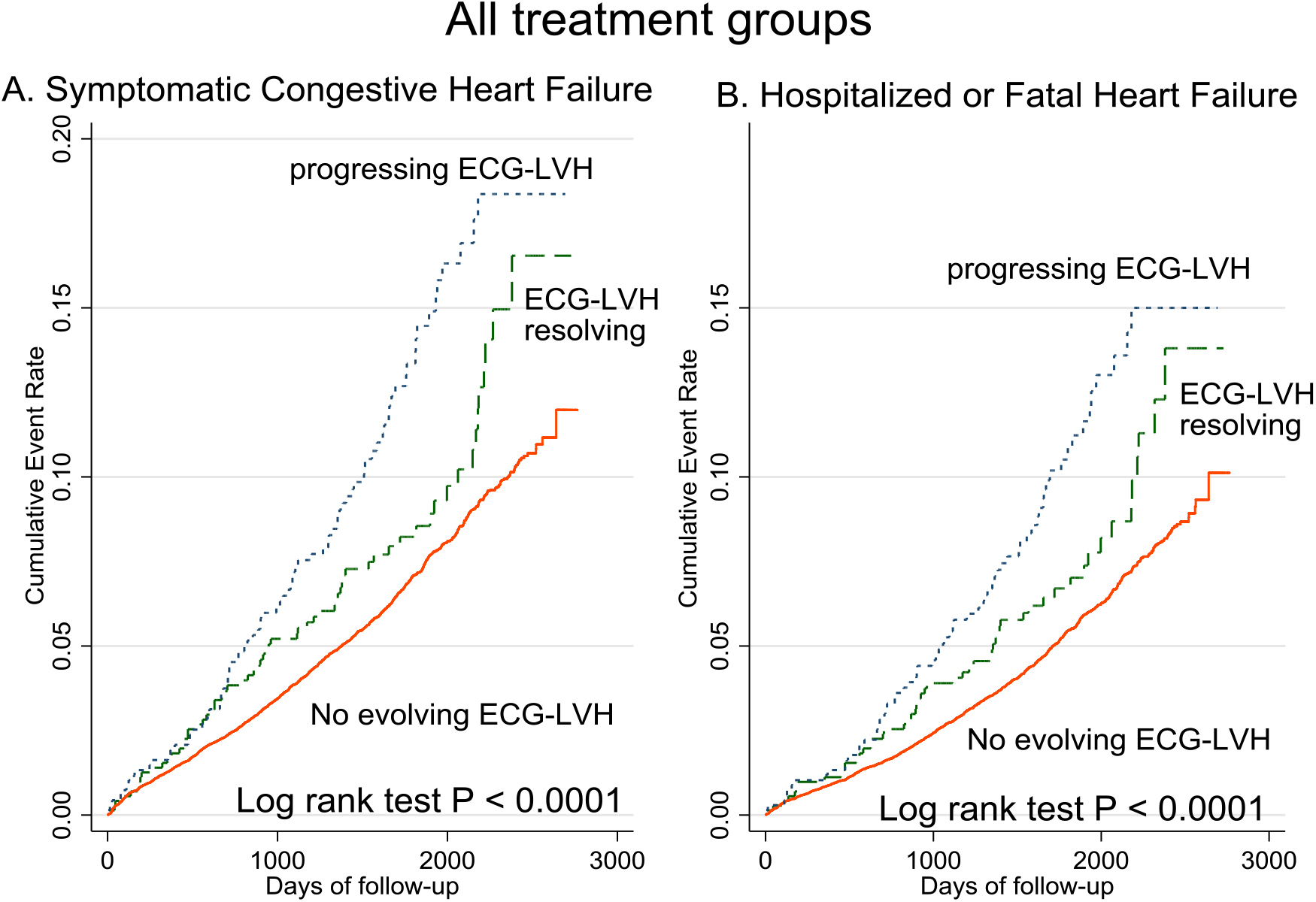
Unadjusted Kaplan-Meier curves for probability of (**A**) incident symptomatic HF and (**B**) hospitalized or fatal HF in all treatment groups ALLHAT participants with evolving ECG-LVH development (blue dotted line), resolution (green dashed line), or without evolving ECG changes (red solid line).

**Figure 5.**
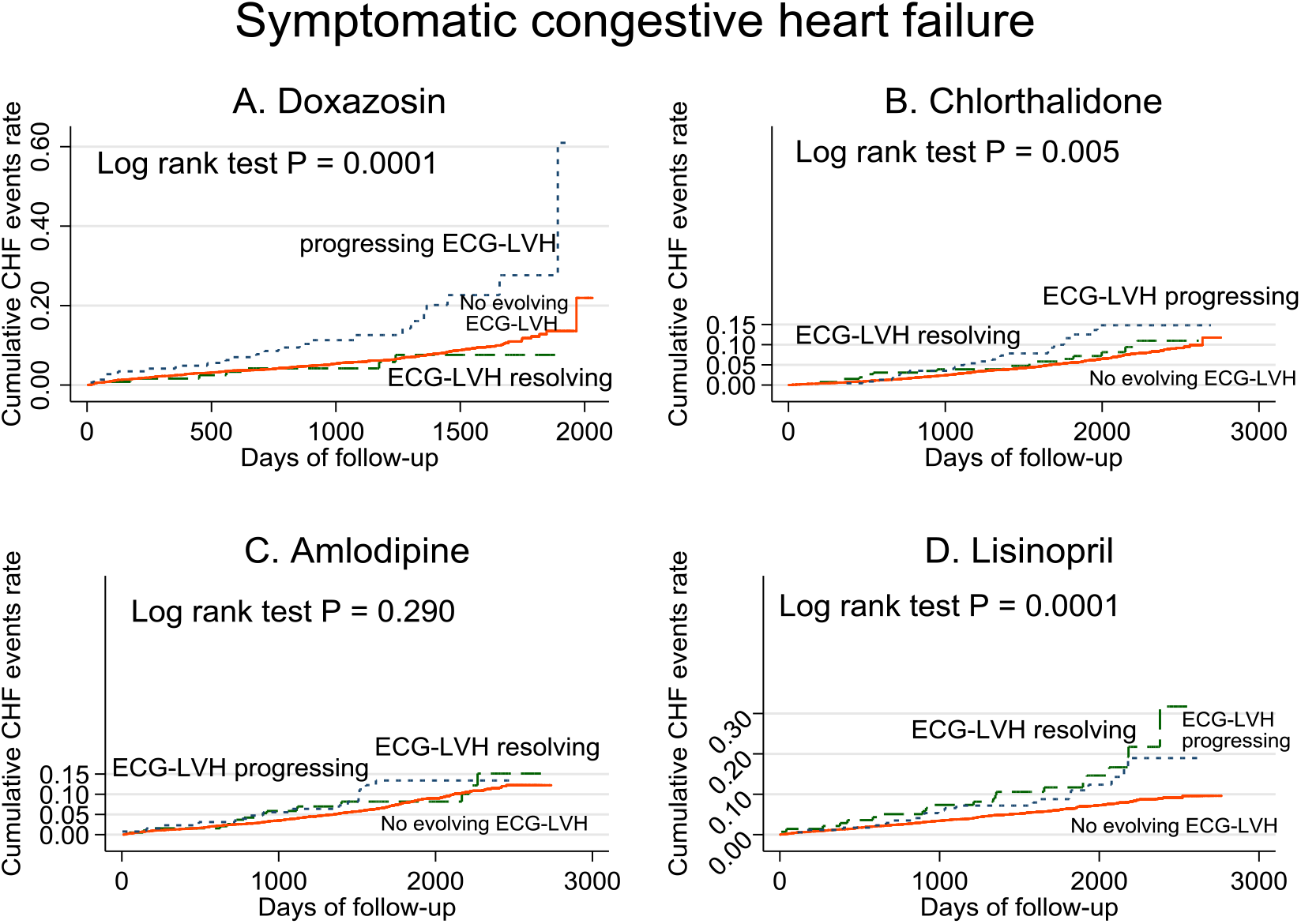
Unadjusted Kaplan-Meier curves for probability of incident symptomatic HF in (**A**) Doxazosin, (**B**) Chlorthalidone, (**C**) Amlodipine, (**D**) Lisinopril treatment groups. Evolving ECG-LVH groups as described in Figure 3 legend.

The association of in-trial BP changes with HF was non-linear (Figure 6). Both large decrease and poor control of BP were associated with incident HF, but large decrease in BP had a stronger effect than poor BP control on both primary and secondary outcomes (Table 4). A similar association of SBP-lowering with incident HF was observed in three out of four treatment groups (Figure 7). In the amlodipine treatment group, SBP change was not associated with incident HF (P_interaction_=0.039; Figure 6). A noticeable U-shaped association of DBP-change with incident symptomatic HF was observed in the amlodipine and chlorthalidone treatment groups (Figure 8), whereas poor DBP control in the lisinopril and doxazosin treatment groups was not associated with incident HF.

**Figure 6.**
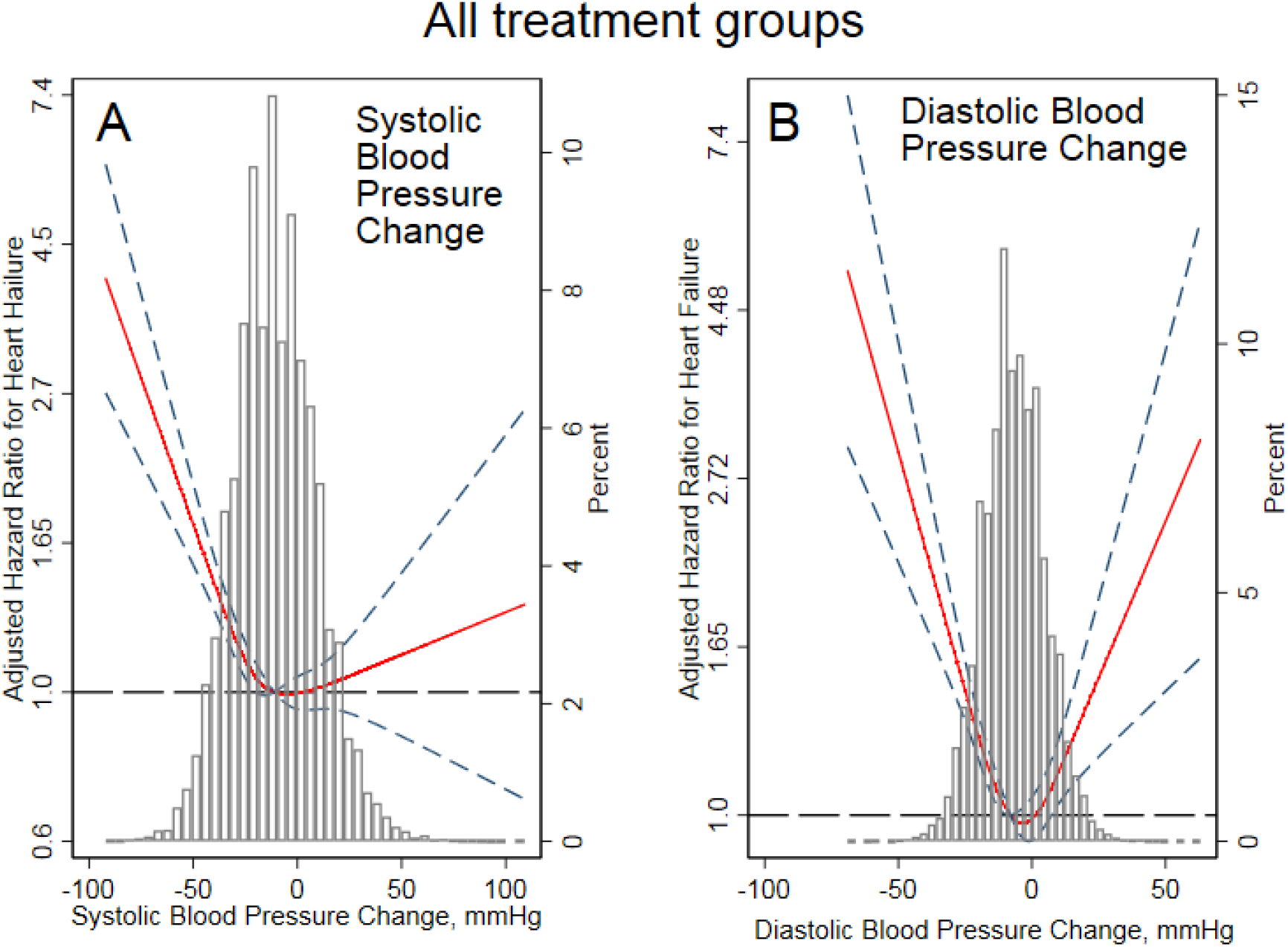
Adjusted (by age, sex, and race/ethnicity) risk of symptomatic congestive HF associated with achieved in-trial greatest SBP and DBP changes, in all participants. Restricted cubic spline with 95% confidence interval show change in hazard ratio (Y-axis) in response to BP change (X-axis). 50^th^ percentile of BP change is selected as the reference.

**Figure 7.**
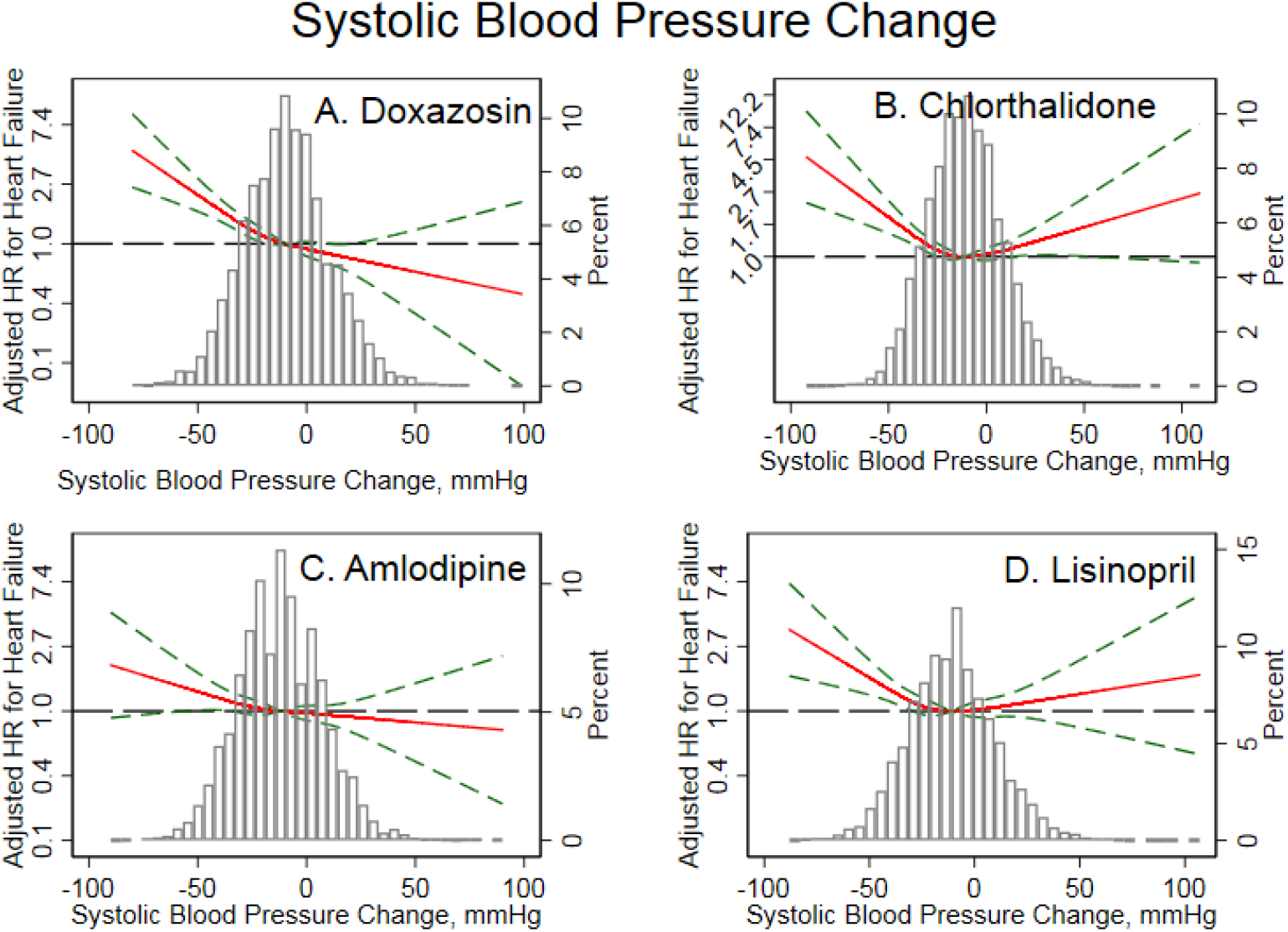
Adjusted risk of symptomatic congestive HF associated with achieved in-trial greatest SBP changes HF in (**A**) Doxazosin, (**B**) Chlorthalidone, (**C**) Amlodipine, (**D**) Lisinopril treatment groups. See Figure 5 legend for details.

**Figure 8.**
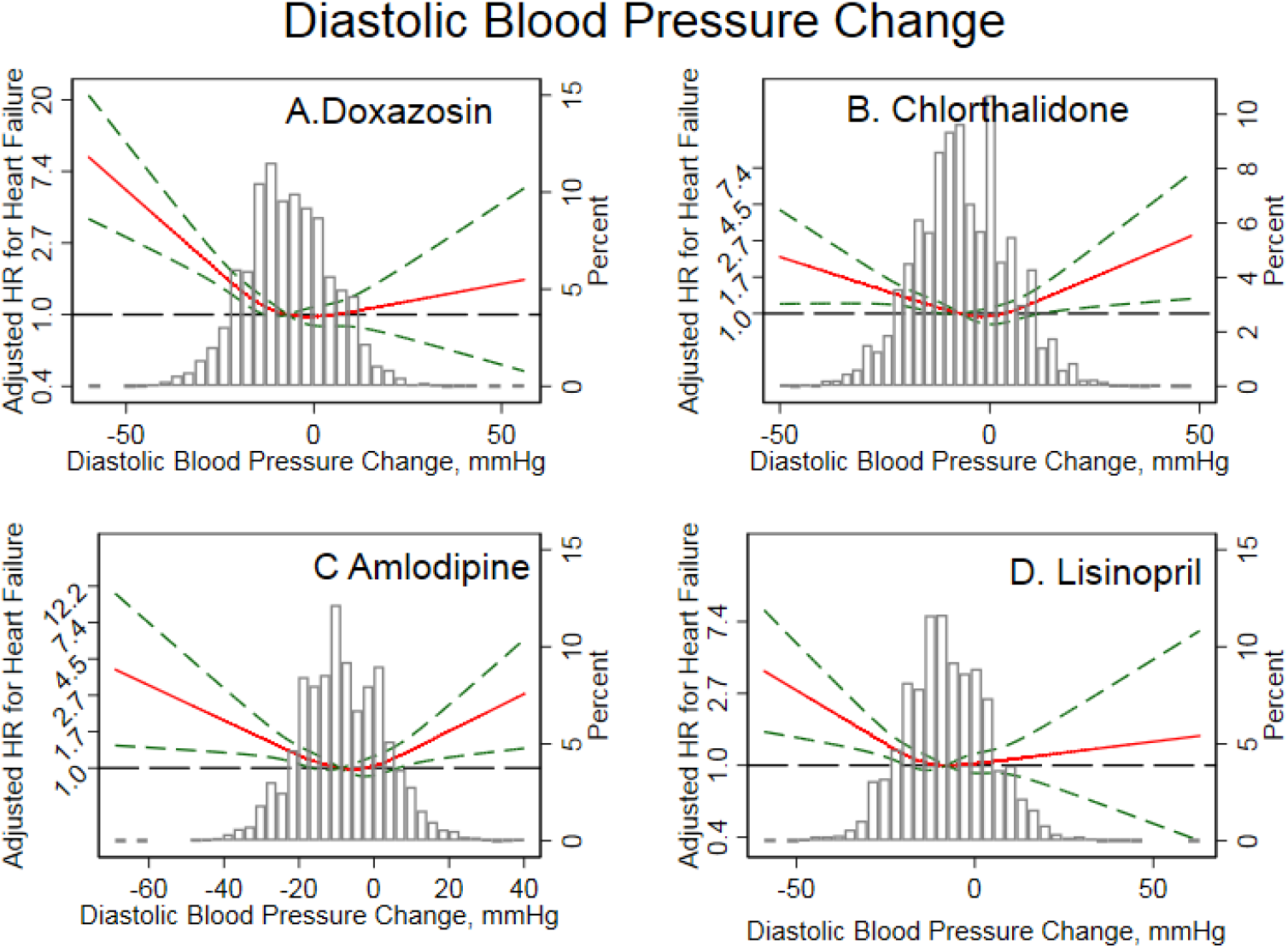
Adjusted risk of symptomatic congestive HF associated with achieved in-trial greatest DBP changes HF in (**A**) Doxazosin, (**B**) Chlorthalidone, (**C**) Amlodipine, (**D**) Lisinopril treatment groups. See Figure 5 legend for details.

### Mediation of HF risk by evolving LVH

In fully adjusted analyses, evolving LVH mediated 4% of the effect of doxazosin on HF (Table 5). Both direct and mediated pathways contributed to the increased HF risk in doxazosin arm. The effect of amlodipine and lisinopril on HF was entirely independent of evolving LVH.

**Table 5.**
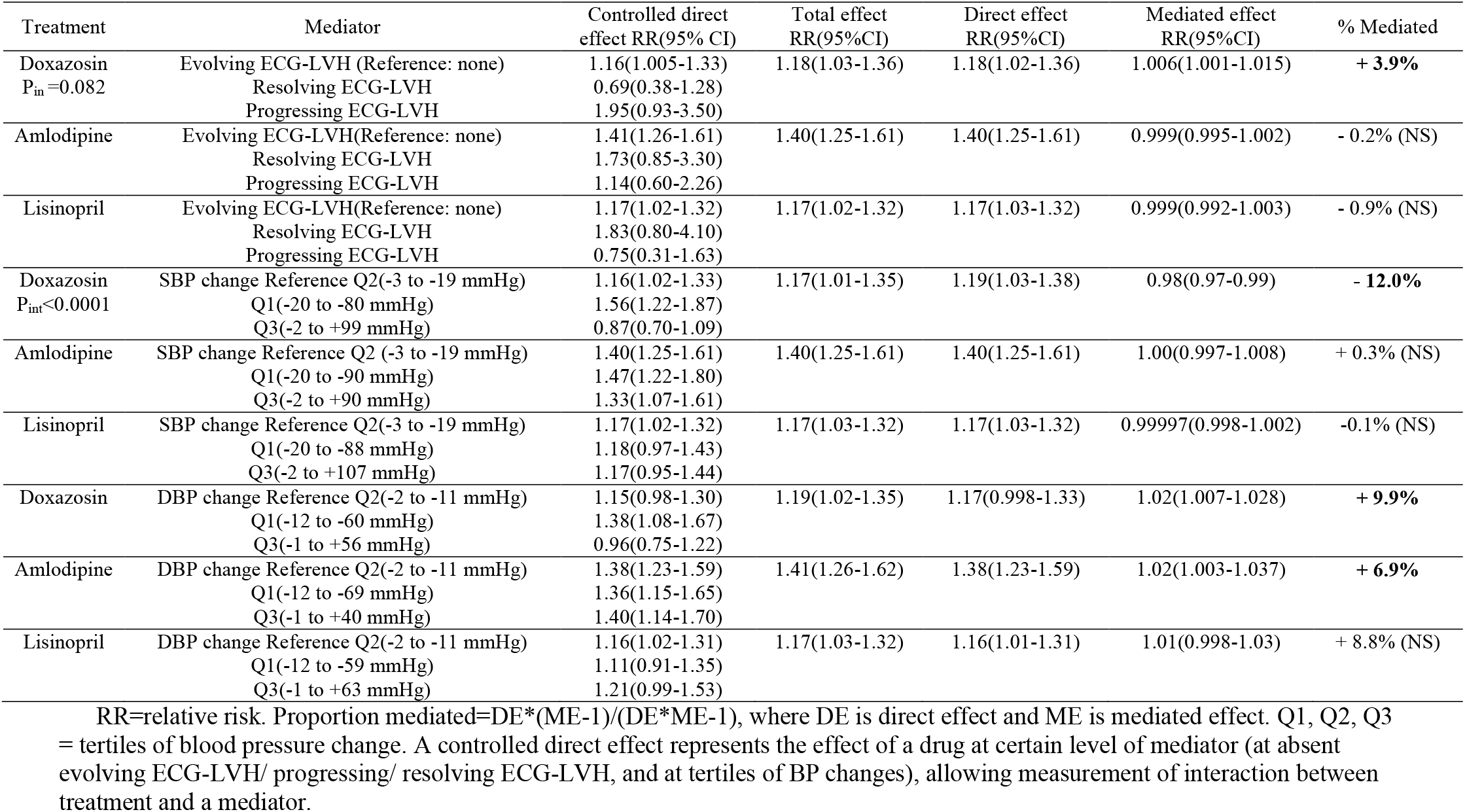
Fully adjusted effect of antihypertensive treatment on incident symptomatic heart failure (total), through evolving ECG-LVH or BP changes (mediated), and independent of BP-lowering or evolving ECG-LVH (direct)

### Mediation of HF risk by dynamic BP changes

After full adjustment for confounders, SBP-lowering mediated 12% of the effect of doxazosin on HF (Table 5). Of note, the direct and mediated effects of doxazosin on HF were in opposite directions: direct effect of doxazosin increased HF risk, whereas SBP-lowering-mediated effect reduced HF risk by 12%. There was significant (P<0.0001) interaction between doxazosin treatment and mediator: SBP-lowering in Q1 and Q2 was associated with increased risk of HF, whereas Q3 SBP change (mean increase 11 mmHg) was protective. The effects of amlodipine and lisinopril on HF were entirely independent of SBP changes.

DBP-lowering mediated 10% of the effect of doxazosin, and 7% of the effect of amlodipine, and 9% of the effect of lisinopril on HF. In fully adjusted analyses (Table 5) mediation of the effect of lisinopril lost statistical significance. Both direct and mediated pathways had the same direction and contributed to the increased HF risk.

Sensitivity analyses with different definitions of BP-lowering provided similar results (Table 6). The fastest SBP-lowering mediated ~13% of the effect of lisinopril on HF.

**Table 6.** Effect of antihypertensive treatment on incident symptomatic heart failure (total), through relative ‘greatest’ BP changes, absolute and relative ‘fastest’ BP changes (mediated), and independent of BP-lowering (direct)

|  | Treatment | Mediator | Total effect<br>RR(95%CI) | Controlled direct<br>effect RR(95% CI) | Direct effect<br>RR(95%CI) | Mediated effect<br>RR(95%CI) | % Mediated |
| --- | --- | --- | --- | --- | --- | --- | --- |
| Relative<br>greatest<br>BP-lowering | Doxazosin | SBP change Q2 | 1.17(0.9996-1.32) | 1.16(1.00-1.32) | 1.19(1.02-1.35) | 0.98(0.97-0.99) | **-13.7%** |
|  | Amlodipine | SBP change Q2 | 1.40(1.25-1.61) | 1.39(1.25-1.61) | 1.40(1.25-1.61) | 1.003(0.998-1.01) | + 1.1% (NS) |
|  | Lisinopril | SBP change Q2 | 1.17(1.03-1.32) | 1.17(1.02-1.32) | 1.17(1.02-1.32) | 1.00(0.997-1.003) | + 0.02% (NS) |
|  | Doxazosin | DBP change Q2 | 1.19(1.02-1.35) | 1.16(0.99-1.31) | 1.17(0.998-1.33) | 1.01(1.006-1.03) | **+ 9.3%** |
|  | Amlodipine | DBP change Q2 | 1.41(1.26-1.62) | 1.38(1.23-1.59) | 1.38(1.23-1.59) | 1.02(1.001-1.04) | **+ 7.0%** |
|  | Lisinopril | DBP change Q2 | 1.17(1.03-1.32) | 1.16(1.02-1.31) | 1.16(1.01-1.31) | 1.01(0.998-1.03) | + 8.8% (NS) |
| Fastest<br>BP-lowering | Doxazosin | SBP change Q2 | 1.17(1.01-1.34) | 1.16(0.998-1.32) | 1.16(0.996-1.32) | 1.008(0.994-1.25) | + 5.6% (NS) |
|  | Amlodipine | SBP change Q2 | 1.40(1.26-1.62) | 1.39(1.25-1.60) | 1.39(1.25-1.60) | 1.006(0.9995-1.01) | + 2.2% (NS) |
|  | Lisinopril | SBP change Q2 | 1.17(1.02-1.32) | 1.15(1.0005-1.29) | 1.15(0.999-1.29) | 1.02(1.009-1.04) | **+13.7%** |
|  | Doxazosin | DBP change Q2 | 1.18(1.01-1.34) | 1.18(1.01-1.34) | 1.18(1.01-1.33) | 1.0005(0.999-1.005) | + 0.3% (NS) |
|  | Amlodipine | DBP change Q2 | 1.41(1.25-1.62) | 1.39(1.24-1.60) | 1.40(1.25-1.61) | 1.004(0.998-1.01) | + 1.2% (NS) |
|  | Lisinopril | DBP change Q2 | 1.17(1.02-1.32) | 1.17(1.02-1.32) | 1.17(1.02-1.31) | 1.001(0.9996-1.006) | + 0.9% (NS) |
| Relative<br>fastest BP-lowering | Doxazosin | SBP change Q2 | 1.17(1.01-1.34) | 1.16(0.998-1.32) | 1.16(0.995-1.33) | 1.01(0.994-1.03) | + 6.3% (NS) |
|  | Amlodipine | SBP change Q2 | 1.40(1.26-1.60) | 1.39(1.25-1.60) | 1.39(1.25-1.60) | 1.007(0.99993-1.02) | + 2.4% (NS) |
|  | Lisinopril | SBP change Q2 | 1.17(1.02-1.32) | 1.15(1.003-1.29) | 1.15(0.999-1.29) | 1.02(1.009-1.035) | **+ 13.4%** |
|  | Doxazosin | DBP change Q2 | 1.18(1.01-1.33) | 1.18(1.01-1.34) | 1.18(1.01-1.33) | 1.0001(0.999-1.003) | + 0.1% (NS) |
|  | Amlodipine | DBP change Q2 | 1.40(1.26-1.62) | 1.40(1.25-1.61) | 1.40(1.25-1.61) | 1.003(0.997-1.01) | + 1.05% (NS) |
|  | Lisinopril | DBP change Q2 | 1.17(1.02-1.32) | 1.17(1.02-1.33) | 1.17(1.02-1.32) | 1.0006(0.9994-1.005) | + 0.4% (NS) |

RR=relative risk. Proportion mediated=DE\*(ME-1)/(DE\*ME-1), where DE is direct effect and ME is mediated effect.
Q2 is a medium tertile of blood pressure changes. A controlled direct effect represents the effect of a drug at the second tertile of BP changes.

## Discussion

The main finding of our study is that the evolving ECG-LVH and BP-lowering explain up to 13% of the HF-preventive effect of diuretic chlorthalidone, as compared to the preventive effect of antihypertensive treatment with the alpha-blocker doxazosin, the ACEi lisinopril, and the CCB amlodipine. This finding highlights the notion of HF as a complex multifactorial condition, and underscores importance of the use of diuretics for HF prevention, which targets mechanisms that are largely independent of BP-lowering and evolving ECG-LVH.

### Heart failure prevention in hypertension

HTN is the major risk factor of HF, associated with 2-3 fold increased HF incidence in observational cohort studies.^17^ However, RCTs HTN treatment is associated with only 20-25% reduction in HF risk^5^. Our study provided consistent findings: BP-lowering mediated only up to 13% effect of antihypertensive medications on incident HF. Such disconnect between a risk factor and effect of its modification is traditionally explained by poor BP control, irreversible damage of the heart over long-time risk exposure, insufficient awareness of HTN, and inadequate assessment of HTN by a single BP measurement. Our study findings suggest that in order to achieve the most effective HF prevention, BP-lowering should not be the only criterion of HTN treatment effectiveness. Moreover, as different antihypertensive treatments have different mediators, different criteria of effectiveness (beyond BP-control) should be developed for each class of antihypertensive drugs.

### Diuretics for HF prevention

Our study showed that mechanisms by which the thiazide diuretic chlorthalidone prevented HF were not restricted to BP-lowering and prevention of LVH. The mechanisms responsible for favorable effect of chlorthalidone on HF prevention in HTN persons are unknown. In addition to BP-lowering, chlorthalidone has pleotropic effects, including improving endothethial function and reducing inflammation and oxidative stress).^18^ Better understanding of the mechanisms behind the effect of chlorthalidone on HF may lead to new drug formulations, specifically targeting HF prevention in patients with HTN.

### Left ventricular hypertrophy and heart failure

Longstanding HTN and LVH can start a devastating cascade that leads to HF via myocyte growth, oxidative stress, and fibrosis.^19^ While antihypertensive drugs have been shown to reduce and even reverse LVH, this study showed that reduction in ECG-LVH increased the risk of HF, as compared to patients who remained free from LVH.

In the current study, evolving LVH mediated only 4% of the effect of doxazosin on HF. Consistent with our findings, previous analysis of Cornell voltage changes during the ALLHAT trial^20^ showed no difference in ECG-LVH development/resolution between the amlodipine, lisinopril, or chlorthalidone treatment arms. There are known limitations of ECG-LVH as a measure of the LV enlargement, as there are more than a dozen ECG-LVH definitions with poor agreement among them.^21^ Differences between LVH measured by ECG vs. LV mass measured by imaging modalities^21^ reflect true differences between the cardiac anatomy and the electrophysiological substrate. ECG-LVH characterizes an abnormal electrophysiological substrate, which is associated with sudden cardiac death and incident HF independent of LV mass and BP control^22–24^. Additional ECG measures of electrophysiological substrate should be considered as potential mediators of antihypertensive treatment effect on HF. For example, sum absolute QRST integral (SAI QRST) was shown associated with HF hospitalization or death in MADIT II study.^25^ Longitudinal changes in global electrical heterogeneity (GEH) were associated with LV dysfunction.^26^ Comprehensive description of electrophysiological substrate beyond evolving LVH (e.g. using SAI QRST and GEH) may improve understanding of mechanisms, responsible for HF development in the setting of HTN.

### Blood pressure lowering and heart failure

Our findings are largely consistent with previous ALLHAT results and conclusions.^27^ Previous analysis of attributable risks due to BP-lowering^28^ concluded that effect of amlodipine on incident HF was BP-independent, whereas BP-lowering only partially explained the effect of lisinopril on HF. In our adjusted mediation analysis, effect of both amlodipine and lisinopril on HF was entirely independent of SBP, whereas DBP-lowering mediated 7% effect of amlodipine and 9% effect of lisinopril. Interestingly, we observed opposite directions of the direct effect of doxazosin (increased HF risk), and SBP-lowering-mediated effect of doxazosin (reduced HF risk by 12%). DBP-lowering mediated 10% effect of doxazosin, and had the same direction with the direct effect of doxazosin. As doxazosin remains a viable HTN treatment option for men with benign prostatic hyperplasia, complex effects of BP-lowering on incident HF should be taken into account for patients on doxazosin. Overall, very modest effect of BP-lowering on incident HF highlights an importance of additional (beyond BP control) biomarkers for assessment of effectiveness of antihypertensive drugs for HF prevention.

### Strengths and Limitations

ALLHAT is the largest RCT of antihypertensive treatment, allowing unbiased mediation analysis, strengthening two major assumptions of mediation analysis. Randomization eliminated exposure-outcome and exposure-mediator confounding. However, limitations of this study should be taken into account. While we adjusted for known common causes of evolving ECG-LVH, BP-lowering, and incident HF, unmeasured confounding can affect this study estimates. ALLHAT enrolled high-risk HTN patients, and results of this study may not be generalizable to a lower-risk populations. In our study, baseline BP displayed moderate correlation with in-trial BP-lowering (Figure 3), which at least partially explained U-shaped association of BP-lowering with incident HF. While we utilized modeling approaches accounting for non-linear associations, it is possible that we under-estimated true effect of BP-lowering on incident HF.

## Acknowledgement

This manuscript was prepared using ALLHAT Research Materials obtained from the NHLBI Biologic Specimen and Data Repository Information Coordinating Center.

## Funding

This work was partially supported by HL118277 (LGT).

## Disclosures

None

